# Individual nucleotide resolution UV cross-linking and immunoprecipitation (iCLIP) to determine protein-RNA interactions

**DOI:** 10.1101/108118

**Authors:** Christopher R. Sibley

**Keywords:** iCLIP, CLIP, RNA-binding protein, RNA, protein-RNA interactions, post-transcriptional regulation

## Abstract

RNA-binding proteins (RBPs) interact with and determine the fate of many cellular RNA transcripts. In doing so they help direct many essential roles in cellular physiology, whilst their perturbed activity can contribute to disease aetiology. In this chapter we detail a functional genomics approach, termed individual nucleotide resolution UV cross-linking and immunoprecipitation (iCLIP), that can determine the interactions of RBPs with their RNA targets in high throughput and at nucleotide resolution. iCLIP achieves this by exploiting UV-induced covalent crosslinks formed between RBPs and their target RNAs to both purify the RBP-RNA complexes under stringent conditions, and to cause reverse transcription stalling that then identifies the direct crosslink sites in the high throughput sequenced cDNA libraries.

## 1. Introduction

The fate of a transcribed RNA is largely determined by interactions with RNA-binding proteins (RBPs) [1,2]. For example, RBPs can regulate the post-transcriptional processing, stability, localization and translation of RNA transcripts as they progresses from transcription to degradation. According to a recent census, ~1542 RBPs [2] are found in humans, with each RBP expected to interact with hundreds-to-thousands of RNA targets. Conversely, individual RNA transcripts can interact with hundreds of RBPs during their lifetime [3,4]. These RBP-RNA interactions occur because the RBPs are recruited to their specific target loci through recognition of specific features (e.g. sequence motifs, secondary/tertiary structures, protein-protein interactions), with these features widely dispersed across the transcriptome. Unsurprisingly, this one-to-many activity means that numerous RBPs have fundamental roles in cell biology [5,6], whilst perturbed activity of several RBPs contributes to various disease aetiologies [1,7–9]. Accordingly, RBPs have attracted considerable interest in recent years, whilst specialized techniques have been established in order to determine the sites of RBP occupancy in a transcriptome-wide manner.

RBP-RNA interactions are primarily studied using variants of the RNA immunoprecipition (RIP) and UV crosslinking and immunoprecipitation (CLIP) techniques. RIP involves the immunoprecipition of a RBP together with its bound RNA that is converted to cDNA then sequenced. Although native immunoprecipitations are most commonly used [10], stability of interactions is in some cases strengthened through use of reversible paraformaldehyde crosslinking. In contrast, CLIP-based approaches initially use UV crosslinking to form an irreversible covalent bond directly between RBPs and their target RNAs.

Unlike the use of paraformaldehyde that can crosslink protein-protein, protein-DNA and protein-RNA interactions, this UV-induced crosslinking is specific to protein-RNA interactions over zero-length distances [11,12]. This allows CLIP to use more stringent biochemical purification of the bound RNAs that reduces the signal-to-noise ratio, and which also helps eliminate non-specific interactions [12–16]. Accordingly, several CLIP protocols have now been developed (Table 1), as they have become the methods of choice for studying protein-RNA interactions.

**Table 1:**
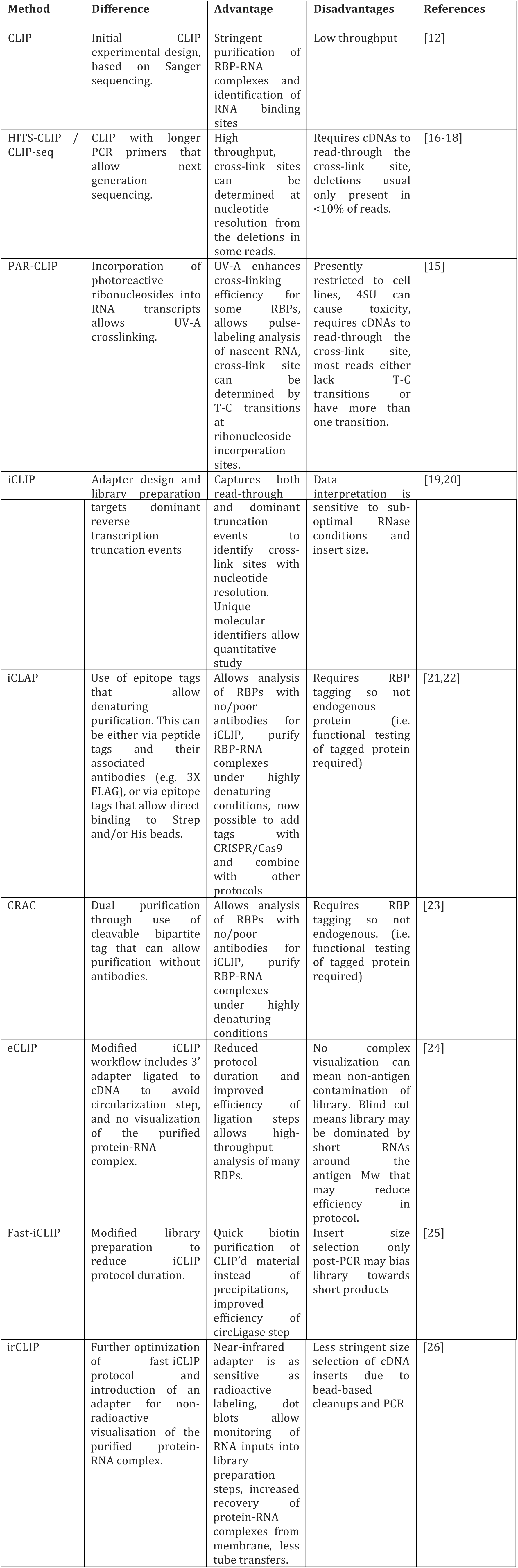
Variants of UV crosslinking and immunoprecipitation

| Method | Difference | Advantage | Disadvantages | References |
| --- | --- | --- | --- | --- |
| CLIP | Initial CLIP experimental design, based on Sanger sequencing. | Stringent purification of RBP-RNA complexes and identification of RNA binding sites | Low throughput | [12] |
| HITS-CLIP / CLIP-seq | CLIP with longer PCR primers that allow next generation sequencing. | High throughput, cross-link sites can be determined at nucleotide resolution from the deletions in some reads. | Requires cDNAs to read-through the cross-link site, deletions usual only present in <10% of reads. | [16-18] |
| PAR-CLIP | Incorporation of photoreactive ribonucleosides into RNA transcripts allows UV-A crosslinking. | UV-A enhances cross-linking efficiency for some RBPs, allows pulse-labeling analysis of nascent RNA, cross-link site can be determined by T-C transitions at ribonucleoside incorporation sites. | Presently restricted to cell lines, 4SU can cause toxicity, requires cDNAs to read-through the cross-link site, most reads either lack T-C transitions or have more than one transition. | [15] |
| iCLIP | Adapter design and library preparation | Captures both read-through | Data interpretation is | [19,20] |
|  | targets dominant reverse transcription truncation events | and dominant truncation events to identify cross-link sites with nucleotide resolution. Unique molecular identifiers allow quantitative study | sensitive to sub-optimal RNase conditions and insert size. |  |
| iCLAP | Use of epitope tags that allow denaturing purification. This can be either via peptide tags and their associated antibodies (e.g. 3X FLAG), or via epitope tags that allow direct binding to Strep and/or His beads. | Allows analysis of RBPs with no/poor antibodies for iCLIP, purify RBP-RNA complexes under highly denaturing conditions, now possible to add tags with CRISPR/Cas9 and combine with other protocols | Requires RBP tagging so not endogenous protein (i.e. functional testing of tagged protein required) | [21,22] |
| CRAC | Dual purification through use of cleavable bipartite tag that can allow purification without antibodies. | Allows analysis of RBPs with no/poor antibodies for iCLIP, purify RBP-RNA complexes under highly denaturing conditions | Requires RBP tagging so not endogenous. (i.e. functional testing of tagged protein required) | [23] |
| eCLIP | Modified iCLIP workflow includes 3' adapter ligated to cDNA to avoid circularization step, and no visualization of the purified protein-RNA complex. | Reduced protocol duration and improved efficiency of ligation steps allows high-throughput analysis of many RBPs. | No complex visualization can mean non-antigen contamination of library. Blind cut means library may be dominated by short RNAs around the antigen Mw that may reduce efficiency in protocol. | [24] |
| Fast-iCLIP | Modified library preparation to reduce iCLIP protocol duration. | Quick biotin purification of CLIP'd material instead of precipitations, improved efficiency of circLigase step | Insert size selection only post-PCR may bias library towards short products | [25] |
| irCLIP | Further optimization of fast-iCLIP protocol and introduction of an adapter for non-radioactive visualisation of the purified protein-RNA complex. | Near-infrared adapter is as sensitive as radioactive labeling, dot blots allow monitoring of RNA inputs into library preparation steps, increased recovery of protein-RNA complexes from membrane, less tube transfers. | Less stringent size selection of cDNA inserts due to bead-based cleanups and PCR | [26] |

In this chapter we discuss the rationale and application of one CLIP method in detail; Individual nucleotide resolution UV-crosslinking and immunoprecipitation (iCLIP)[13,19]. Like other CLIP approaches, iCLIP involves cross-linking RBPs to their RNA targets, immunoprecipitating the RBP-of-interest, digesting away the RBP, and converting the bound RNA into a cDNA library that can be high-throughput sequenced. The advantage of iCLIP over other methods is that it specifically exploits the fact that ~80-100% of cDNAs produced during the reverse transcription step of the protocol truncate at the protein-RNA cross-link site (Figure 1A). This truncation has been both experimentally and computationally validated [27,20], and implies that CLIP protocols requiring crosslink read-through by the reverse transcriptase (e.g. HITS-CLIP, CLIP-Seq, PAR-CLIP) are losing information during library production. In contrast, by using an adapter configuration that captures both truncation events and read-through events (Figure 1B), iCLIP allows identification of the cross-link site with nucleotide resolution and permits quantitative assessment of RBP-binding activity [19,28]. The eCLIP protocol additionally exploits this truncation event using a different adapter configuration, but fundamentally differs from iCLIP in its absence of a protein-RNA complex visualization step [24]. We always recommend complex visualization to help identify or exclude contaminating complexes contributing to the signal, reveal dimers/trimers that may be of interest, allow accurate isolation of RNAs that are of suitable length for library construction, and to provide an additional purification of free RNA that may stick to the beads used in the immunoprecipitation.

**Figure 1.**
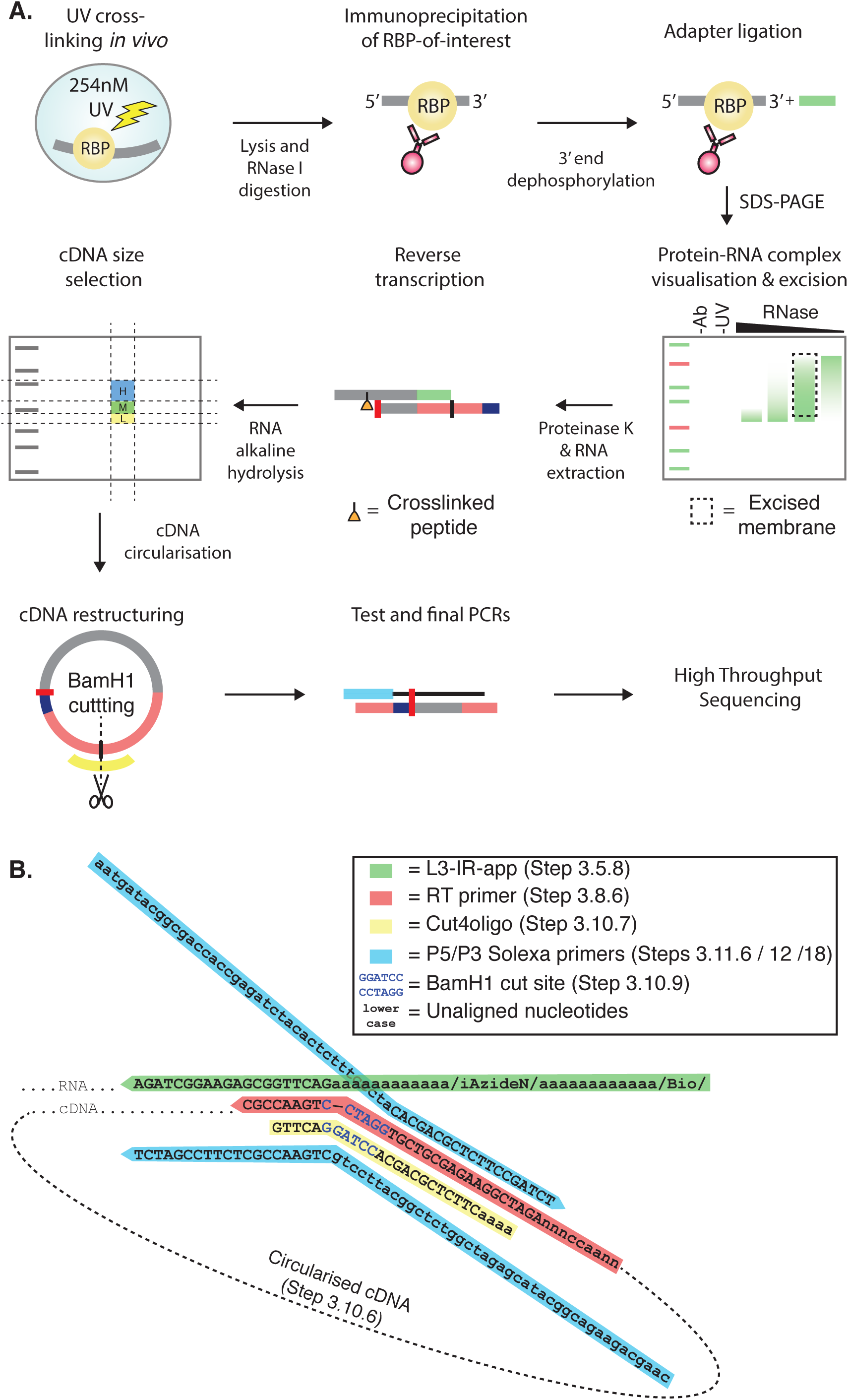
The iCLIP protocol. **A)** The iCLIP protocol begins by cross-linking samples in order to covalently bind RBPs and their RNA targets together. Samples are then lysed before a controlled RNase I digestion is carried out in order to shorten cross-linked RNA fragments to lengths compatible with RNA sequencing. At this point the RBP-of-interest is immunoprecipitated together with its bound cargo. Due to the presence of the covalent bond this purification can stringent to remove non-specific interactions. This can include contaminating RBPs in complex with the RBP-of-interest, RBPs which non-specifically bind to the beads, or RNAs which stick to the beads. Once on the beads, the exposed RNA termini are manipulated to allow a universal adapter to be ligated to the 3’ ends. In addition to providing a specified sequence with which to carry out reverse transcription during library production (Figure 1B), the adapter has a fluorophore attached which allows analysis of the protein-RNA complexes following SDS-PAGE and transfer to nitrocellulose membrane to remove non-crosslinked RNA (Figure 2A). Protein-RNA complexes are then purified from the membrane using a cutting-mask made from the fluorescent image, before the protein is removed by proteinase K digestion and RNA extracted ahead of cDNA library preparation. Library preparation begins with reverse transcription using bipartite adapters which are complementary to the ligated adapter at their 3’ end (Figure 1B). The remainder of the primer completes the sequence of a 3’ solexa sequencing primer, and also contains a juxtaposed 5’ solexa sequence in the opposing orientation followed by experimental barcodes (Figure 1B). The reverse transcription reaction will truncate at the cross-link site, which still retains a covalently bound short polypeptide, in approximately 80-100% of reactions. Following cDNA size selection and primer removal using a cDNA cutting mask (Figure 3) from a denaturing gel, the 5’ solexa sequence and barcodes of the reverse transcription primer are ligated to the 3’ end of the cDNA in a circularization reaction. The 3’ cDNA end corresponds either to the truncation site (80-100%) or read-through sequence (0-20%), thereby capturing all CLIP events unlike other methods (Table 1). These can be distinguished bioinformatically following sequencing. A BamH1 digestion of the circularized cDNA at a site in between the 5’ and 3’ solexa primers results in a linear cDNA with 5’ and 3’ solexa sequences appropriately located at either termini to permit PCR amplification of the library ahead of quantification and next generation sequencing (Figure 4). Importantly, random barcodes contained within the reverse transcription primer barcode region allow PCR duplicates to be filtered out during computational analysis to ensure quantitative nature of the approach is maintained. **B)** Adapter and oligonucleotide alignments for the iCLIP protocol. Colours are matched to the adapters and oligonucleotides shown in Figure 1A.

The iCLIP protocol has been extensively optimized in recent years, and we refer the reader to two closely related publications from the Ule [13] and Konig [29] research groups that will assist with experimental design alongside the current chapter. In the described protocol we additionally incorporate the use of a non-radioactive adapter that will permit a broader use of the iCLIP approach than the previous radioactivity-dependent method. This adapter was recently introduced in the related irCLIP method from the Khavari group, and has potential to allow accurate quantification of RNA inputs at different steps throughout library preparation [26]. These additional benefits are not described in this chapter and the reader is directed towards the irCLIP manuscript. Instead, here the adapter is used simply as a substitute of the traditional iCLIP adapter described previously.

The iCLIP approach is a multi-step process and care must be taken at each in order to ensure protocol success and library fidelity. The appropriate use of controls can allow the protocol to be troubleshooted in real time. However, section 4 provides additional tricks and troubleshooting tips to assist further.

## 2. Materials

### 2.1. Required equipment and consumables

1. 254nm UV crosslinker
2. PAGE Electrophoresis module (e.g. ThermoFisher Xcell II)
3. PAGE Transfer module (e.g. ThermoFisher Xcell II)
4. Thermocycler
5. qPCR machine
6. 1.5 ml / 2ml tube thermomixer
7. 1.5 ml centrifuge
8. Vacuum pump
9. Sonicator
10. Magnetic rack
11. Acetate printing film
12. Printer
13. Near infrared imager (see **Note 1**)
14. Cell scrapers
15. 15ml centrifuge tubes
16. 2 ml microcentrifuge tubes
17. 1.5 ml microcentrifuge tubes
18. Low-binding 0.5 ml microcentrifuge tubes (e.g. Thermofisher AM12350)
19. Low-binding 1.5 ml microcentrifuge tubes (e.g. Thermofisher AM12450)
20. PCR tubes
21. qPCR tubes/microplates
22. Sterile filters
23. Protran 0.45 nitrocellulose membrane
24. Whatman filter paper
25. Razor blades
26. 30G syringe needles
27. 16G syringe needles
28. Phase-lock heavy columns
29. Costar-X filter spin columns (e.g. VWR International)
30. Proteus Clarification columns (e.g. Generon)
31. Glass pre-filters

### 2.2. Required reagents and solutions

1. 1X PBS
2. Tris-HCl pH 7.4
3. Tris-HCl pH 6.5
4. Tris-HCl pH 7.8
5. NaCl solution
6. Igepal CA-630
7. SDS
8. Sodium deoxycholate
9. Urea
10. EDTA pH 8.0
11. MgCl_2_
12. Tween-20
13. Dithiothreitol
14. PEG400
15. LDS-4X sample buffer
16. Methanol
17. 20X MOPS-SDS running buffer
18. 20X Transfer buffer
19. Neutral Phenol:Chloroform
20. TE buffer pH 7.0
21. TE buffer pH 8.0
22. 3M sodium acetate, pH 5.5
23. 80% and 100% Ethanol (molecular biology grade)
24. 1M NaOH
25. HEPES pH 7.3
26. 2X TBE-UREA loading buffer
27. TBE buffer
28. Magnetic protein G/A beads
29. RBP antibodies
30. Anti-hnRNP C (positive control) (e.g. Santa Cruz sc 32308)
31. Protease inhibitor cocktail
32. Anti-RNase antibody (e.g. Thermo Fisher AM 2690)
33. RNase I (e.g. Thermo Fisher)
34. Turbo DNase (e.g. Thermo Fisher)
35. T4 PNK (e.g. NEB)
36. RNase inhibitor (e.g. Promgea RNasin)
37. T4 RNA ligase I
38. Near infrared protein marker (e.g. Li-Cor Chameleon)
39. Antioxidant (e.g. Life Technologies NP0005)
40. Reducing agent (e.g. Life Technologies NP0004)
41. Proteinase K (e.g. NEB)
42. Glycoblue
43. 10 mM dNTPs
44. Reverse transcriptase (e.g. ThermoFisher Superscript III)
45. Low molecular DNA weight marker (e.g. NEB)
46. Sybr safe
47. Circligase II, MnCl2 and 10X buffer
48. Fast Digest BamH1
49. PCR mastermix (e.g. Thermo Fisher Accuprime supermix I)
50. Illumina qPCR library quantification kit (e.g. Kapa biosystems)
51. 4-12% protein denaturing pre-cast gels (e.g. Thermo Fisher NuPAGE) (see **Note 2**)
52. 6% TBE-UREA pre-cast gels (e.g. Thermo Fisher)
53. 6% TBE pre-cast gels (e.g. Thermo Fisher)

### 2.3. Required oligonucleotides

1. L3-IR-app adapter: /5Phos/AGATCGGAAGAGCGGTTCAGAAAAAAAAAAAA/iAzideN/AAAAAA AAAAAA/3Bio/ Adapter requires adenylation then click chemistry conjugation of the IRDye 800CW DBCO infrared dye (Li-Cor) before use.
2. RT-primers: /5Phos/NN<u>AACC</u>NNNAGATCGGAAGAGCGTCGTGgatcCTGAACCGC underlined nucleotides can be replaced with different index barcodes e.g. RT1: AACC, RT2: ACAA, RT3: ATTG, RT4: AGGT, RT6: CCGG, RT7: CTAA, RT8: CATT
3. Cut4oligo: 5’GTTCAGGATCCACGACGCTCTTCaaaa
4. P5 primer: 5’AATGATACGGCGACCACCGAGATCTACACTCTTTCCCTACACGACGCTCTTCCGATCT
5. P3primer: 5’CAAGCAGAAGACGGCATACGAGATCGGTCTCGGCATTCCTGCTGAACCGCTCTTCCGATCT

### 2.4. Buffers

1. Lysis Buffer: 50 mM Tris-HCl, pH 7.4, 100 mM NaCl, 1% Igepal CA-630, 0.1% SDS, 0.5% sodium deoxycholate. Sterile filter and store at 4 °C
2. High-salt Wash: 50 mM Tris-HCl, pH 7.4, 1 M NaCl, 1 mM EDT, 1% Igepal CA-630, 0.1% SDS, 0.5% sodium deoxycholate. Sterile filter and store at 4 °C.
3. PNK Buffer: 20 mM Tris-HCl, pH 7.4, 10 mM MgCl_2_, 0.2% Tween-20. Sterile filter and store at 4 °C.
4. 5x PNK pH 6.5 Buffer: 350 mM Tris-HCl, pH 6.5, 50 mM MgCl_2_, 5 mM dithiothreitol. Use nuclease-free H_2_O. Dispense to 50 μl aliquots and store at −20°C. Only use aliquots once.
5. 4x Ligation Buffer: 200 mM Tris-HCl, pH 7.8, 40 mM MgCl_2_, 4 mM dithiothreitol. Use nuclease-free H_2_O. Dispense to 50 μl aliquots and store at −20°C. Only use aliquots once.
6. PK Buffer: 100 mM Tris-HCl, pH 7.4, 50 mM NaCl, 10 mM EDTA. Sterile filter and store at 4 °C.
7. PK Buffer + 7 M Urea: 100 mM Tris-HCl, pH 7.4, 50 mM NaCl, 10 mM EDTA, 7 M Urea. Sterile filter and store at 4 °C.
8. 1x NuPAGE MOPS-SDS buffer: Add 25 ml of 20X MOPS-SDS buffer to 475 ml of water
9. 1x NuPAGE loading buffer (per sample): 5 μl 4x NuPAGE Loading Buffer, 2 μl sample reducing reagent, 13 μl nuclease-free H_2_O

### 2.5 Reaction mixes

1. PNK de-phosphorylation mix (per sample): 15 μl nuclease-free H_2_O, 4 μl of 5x PNK pH 6.5 Buffer, 0.5 μl PNK, 0.5 μl RNAsin
2. Ligation mix (add reagents in the indicated order): 8 μl nuclease-free H_2_O, 5 μl of 4x ligation buffer, 0.5 μl T4 RNA ligase, 0.5 μl RNAsin, 1.5 μl 1 μM L3-IR-app oligo, 4 μl PEG 400
3. Reverse transcription mix (per sample): 7 μl nuclease-free H_2_O, 4 μl 5x RT buffer, 1 μl 0.1 M DTT, 0.5 μl RNasin, 0.5 μl Superscript III.
4. Circularisation mix (per sample): 6.5 μl nuclease-free H_2_O, 0.8 μl 10x CircLigase II buffer, 0.4 μl 50 mM MnCl2, 03 μl CircLigase II.
5. Cut oligo mix(per sample): 25 μl nuclease-free H_2_O, 4 μl FastDigest buffer, 1 μl 10 μM cut oligo
6. Test-PCR mix (per sample): 3.75 μl ddH_2_O, 5 μl AccuPrime Supermix 1, 0.25 μl 10 μM P5/P3 solexa primer mix
7. Final-PCR mix (per sample): 9 μl ddH_2_O, 20 μl AccuPrime Supermix 1, 1 μl 10 μM P5/P3 solexa primer mix
8. Universal qPCR Master mix (per sample): 12.4 μl qPCR Master mix (Primer Premix and ROX High/Low should be added to Master Mix prior to first use as per manufacturers instructions), 3.6 μl ddH_2_O

## 3. Methods

### 3.1 Sample collection

1. Remove media from cells growing at 80-90% confluency on a 10cm dish and replace with 6 mls of ice cold PBS (see **Note 3**). **Important: Samples should remain at 4°C from this point forward unless indicated**. **Optional: To test if the signal is dependent on expected antigen and not a contaminant RBP, use a control cell-line in which the RBP of interested is knocked-down or knocked-out (see Note 4)**.
2. Place plate on ice-filled tray which has dimensions suitable for UV crosslinker (see **Note 5**).
3. Ensure cell culture dish lids are removed before irradiating cells at 150 mJ/cm^2^ at 254 nm. **Important: Remember to proceed with some plates to step 4 with no crosslinking. These will be used as a no UV control (see Note 6)**.
4. Immediately harvest cells through gentle use of a cell scraper. Cells should be aliquoted into 2 ml samples (see **Note 7**).
5. Spin cell suspensions at 376x g for 1 min at 4 °C to pellet cells.
6. Remove supernatant and snap freeze cells on dry ice. Store cell pellets at −80 °C until use.

### 3.2 Bead preparation

1. Add 100 μl of protein G or protein A magnetic beads to a microcentrifuge tube (see **Note 8**).
2. Place microcentrifuge tube on a magnetic rack, remove supernatant, then wash beads twice in 900 μl of lysis buffer. Beads should be resuspended by rotation for each wash (see **Note 9**).
3. Resuspend in 100 μl per sample of lysis buffer.
4. Transfer 100 μl of bead suspension to a second microcentrifuge tube to act as no antibody control in order to confirm that the signal is dependent on antigen and not unspecific RBPs or RNA binding to beads (see **Note 10**).
5. To the first microcentrifuge tube add 2-10 μg antibody per sample (see **Note 11**).
6. Rotate tubes in a cold room for 30-60 mins whilst proceeding with sample preparation (section 3.3).
7. When lysate is ready wash beads in 1 x high salt wash and 2 x lysis buffer. After final wash resuspend beads in 100 μl lysis buffer per sample. Proceed to step 4 of section 3.4.

### 3.3 Sample preparation

1. Prepare 2.1 ml of ice cold lysis buffer per sample by adding 21 μl (1:100) proteinase inhibitor cocktail. For starting material with high RNase activity add an additional 2.1 μl anti-RNase per sample.
2. Re-suspend cell pellets in 1ml of prepared lysis buffer and place on ice. **Recommended: At this stage the protein content of the lysates can be assessed with a Bradford assay. Samples using single pellets should be ~ 2 mg / ml and standardised by removing lysate from samples with excess. Removed volumes of lysate should be replaced with fresh lysis buffer to ensure identical 1 ml volumes.**
3. Prepare a 1/1000 dilution of RNase I in 500 μl of ice cold lysis buffer. This dilution is used for low RNase samples (see **Note 12**).
4. Prepare a 1/10 dilution of RNase I in 15 μl of ice cold lysis buffer for high RNase condition (see **Note 13**).
5. Add 10 μl of low RNase dilution to all lysates together with 2 μl Turbo DNase. To high RNase sample add additional 10 μl of high RNase dilution.
6. Incubate samples for exactly 3 min at 37°C whilst shaking at 1100 rpm. After incubation immediately transfer samples to ice for 3 min.
7. Centrifuge lysates at >18,000x g for 10 min at 4 °C and transfer lysate to new 1.5 ml microcentrifuge tube. Take care not to disturb pelleted debris during transfer.
8. Load 500 μl of lysate per sample into a Proteus Clarification spin column and spin at >18,000x g for 1 min. Transfer flow-through to a new 2 ml microcentrifuge tube. Repeat step with remaining lysate and combine accordingly.
9. Add 1 ml of prepared ice cold lysis buffer and place on ice until beads are washed and ready for immunoprecipitation. **Optional: At this stage 20 μl of lysate can be retained as a pre-immunoprecipitation sample and used to evaluate immunoprecipitation efficiency through western analysis.**

### 3.4 Immunoprecipitation

1. Ensure beads are uniformly resuspended before adding 100 μl of beads to cell lysates.
2. Rotate beads for 1 hr at 4°C in cold room. **Optional: Once beads have been separated from lysate on a magnetic rack, 20 μl of lysate can be retained as a post-immunoprecipitation sample. This can be used with previously collected sample to evaluate immunoprecipitation efficiency through western analysis.**
3. Discard supernatant and wash beads in 2 x high salt wash. Rotate the second wash for 5 mins in the cold room.
4. Wash beads 2 x PNK wash buffer and re-suspend in 1 ml PNK wash buffer until ready to proceed with next step.

### 3.5 Adapter ligation

1. Remove PNK wash buffer from beads and remove microcentrifuge tubes from magnet for 30 secs. Return microcentrifuge tubes to magnet and remove any small volumes of buffer still retained (see **Note 14**).
2. Remove beads from magnet and resuspend in 20 μl of de-phosphorylation mix.
3. Incubate samples for exactly 20 min at 37°C whilst shacking at 1100 rpm.
4. Wash with 1 x PNK buffer.
5. Wash with 1 x high salt wash buffer. Rotate the wash for 5 mins in the cold room.
6. Wash with 2 x PNK buffer and leave in final wash until ready for ligation.
7. Remove PNK buffer from beads and remove microcentrifuge tubes from magnet for 30 secs. Return microcentrifuge tubes to magnet and remove any small volumes of buffer still retained (see **Note 14**).
8. Remove beads from magnet and resuspend in 20 μl of ligation mix.
9. Incubate samples overnight at 16°C whilst shaking at 1100 rpm.
10. Wash with 1 x PNK buffer.
11. Wash with 2 x high salt wash buffer. Rotate the first wash for 5 mins in the cold room (see **Note 15**).
12. Wash with 1 x PNK buffer and transfer to new 1.5 ml microcentrifuge tube.
13. Wash with 1 x PNK buffer and leave in final wash until ready for SDS-PAGE.

### 3.6 Protein-RNA complex visualisation

1. Assemble the XCell III gel system according to manufacturers instructions using a 4-12% NuPAGE Bis-Tris gel (see **Note 2**) and 1x MOPS-SDS buffer.
2. Add 500 μl of antioxidant to the upper chamber (see **Note 16**).
3. Remove PNK buffer from beads and remove microcentrifuge tubes from magnet for 30 secs. Return microcentrifuge tubes to magnet and remove any small volumes of buffer still retained (see **Note 14**).
4. Re-suspend beads in 20 μl of 1X NuPage sample loading buffer.
5. Incubate samples at 80 °C for 5 mins to dissociate sample from beads. The chameleon protein ladder does not need to be heated.
6. Spin down any precipitation of the samples using a desktop microcentrifuge then place on magnet to separate beads.
7. Load 20 μl of sample supernatant to gel using gel loading tips, and 5 μl of chameleon ladder. It is suggested that gaps are left between samples to facilitate extraction of protein-RNA complexes from the membrane.
8. Run the gel for the 50 mins at 180 V
9. Prepare fresh transfer buffer by adding 25 ml of 20X transfer buffer and 50 ml of methanol to 425 ml of water
10. Following SDS-PAGE remove gel cassette and carefully open. Assemble blotting ‘sandwich’ and XCell III blotting module as per manufacturers instructions.
11. Transfer protein-RNA complexes for 1.5 hr at 30 V
12. Remove nitrocellulose transfer membrane and transfer to a light protected PBS containing box.
13. Visualise protein-RNA complexes using appropriate fluorescent imager and return to PBS until protein-RNA complex isolation (see Figure 2A).

**Figure 2:**
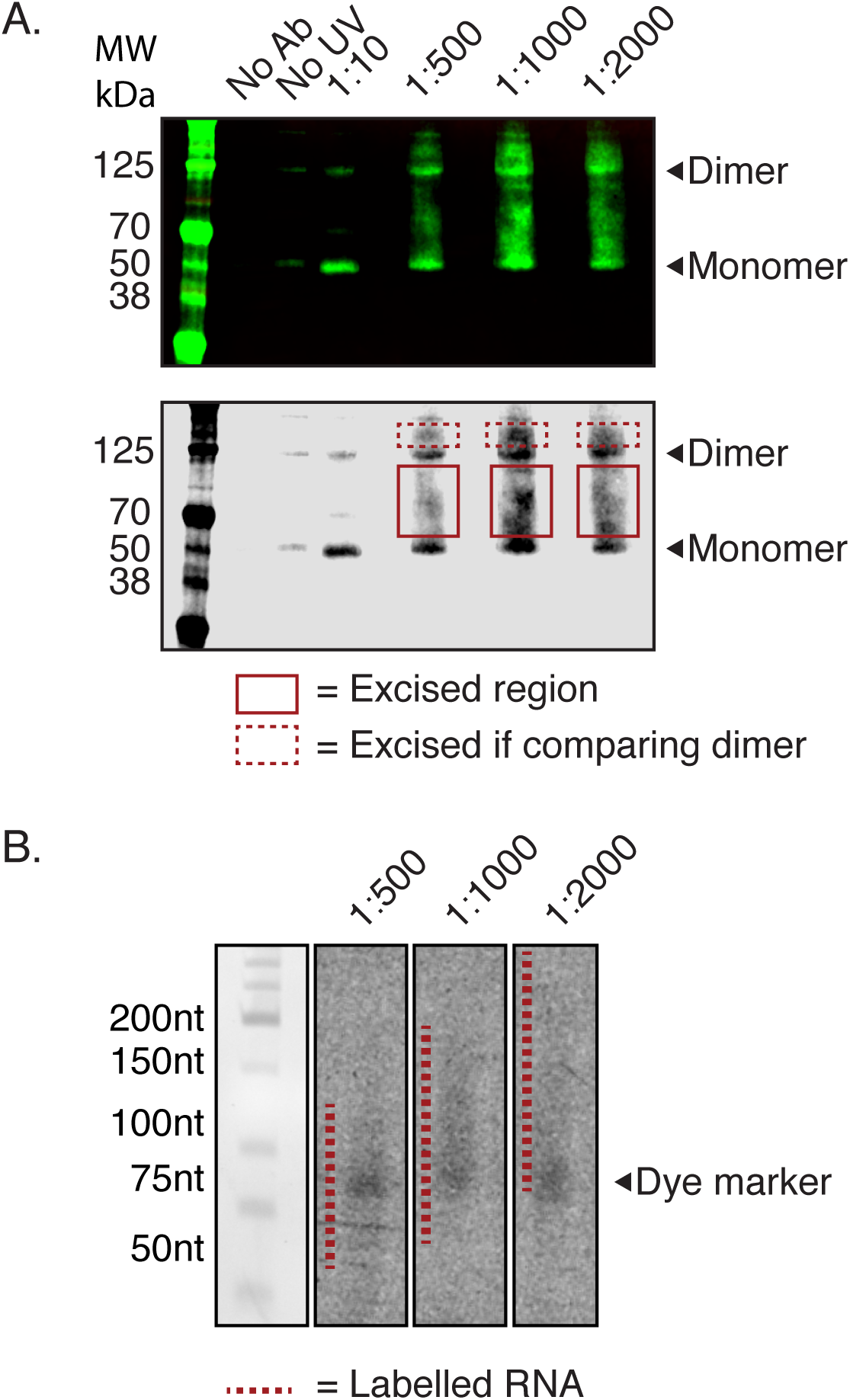
Protein-RNA complex visualisation and RNase digestion analysis. **A)** Fluorescent adapter labelled RNA-protein complexes visualised following SDS-PAGE. Upper panels shows expected outcome on Li-cor imager, lower panel shows grayscale version used to aid complex excisions from the membrane (see step 14 section 3.6). Samples include no UV and no antibody negative controls, and a gradient of RNase I conditions from which suitable digestions patterns can be determined. The immunoprecipitated RBP is hnRNP C, and the arrowhead indicates both monomers and dimers particularly discernable in the high RNase condition. Note that once optimal digestion conditions have been determined for RNase I on the desired sample batch, only the high RNase and determined concentration (in this case 1:1000) need to be carried out for each new iCLIP experiment alongside the no UV and no antibody negative controls. The solid red markers indicate regions excised from the membrane using a cutting mask made from the gel image. The dotted red markers indicate region that would be cut if comparisons between monomer and dimer were to be made. **B)** RNA size distributions from the different regions and different RNase concentrations indicated in cut regions from Figure 2A. Extracted RNA was denatured and run for 40 minutes at 180V on a 6% TBE-UREA gel. Dotted red line indicates detectable size distributions of each RNase condition. Signal intensity can be limited relative to alternative radiolabelling methods due to maximum exposure time limits on some imaging machines.
14. Print image at 100% scale on acetate film. It is recommended that a grey scale image be printed to facilitate alignment of ladder markers between the film and the membrane (see Figure 2A, lower panel).

### 3.7 Protein-RNA complex isolation and RNA purification

1. Make a cutting mask to guide protein-RNA complex excision by drawing a box around protein-RNA signal that starts just above the Mw of the RBP-of-interest (see **Note 17**). The high RNase condition can be used to assist accurate assessment of the antigens Mw. Cut out these boxes from the acetate film to create windows that locate protein-RNA signal (Figure 2A).
2. Remove transfer membrane from PBS and wrap in saran wrap. Secure firmly to fixed cutting surface.
3. Align cutting mask with membrane by matching protein ladder markers. Use windows to guide removal of nitrocellulose membrane segments containing protein-RNA complex. Remove co-excised saran wrap and cut membrane sections into small pieces. Transfer to 1.5 ml microcentrifuge tube with the assistance of a 30G syringe needle (see **Note 18**). **Recommended: The cut membrane can be re-analysed to assess cutting accuracy.**
4. Prepare proteinase K digest mix by adding 10 μl of proteinase K to 200 μl of PK buffer per sample
5. Add 200 μl of proteinase K digest mix to nitrocellulose membrane pieces and incubate for 20 mins at 37°C whilst shaking at 1100 rpm.
6. Add 200 μl of PK-urea buffer and incubate for an additional 20 mins at 37°C whilst shaking at 1100 rpm.
7. Transfer supernatant to Phase Lock Heavy gel columns together with 400 μl of neutral phenol-chloroform.
8. Incubate for 5 mins at 30°C whilst shaking at 1100 rpm.
9. Separate phases by centrifuging at >18,000g at room temperature for 5 mins.
10. Taking care not to touch the gel matrix, transfer the aqueous upper phase to a new low-binding 1.5 ml microcentrifuge tube.
11. Spin at >18,000x g for 1 min then transfer to a new low-binding 1.5 ml microcentrifuge tube.
12. Purify RNA by adding 0.75 μl of Glycoblue and 40 μl of 3M sodium acetate pH 5.5 then mixing. Add 1ml of ice-cold 100% ethanol, mix again, then precipitate overnight at −20°C.

### 3.8 Reverse Transcription

1. Spin samples for 20 mins at 4°C and >18,000x g.
2. Remove supernatant leaving ~50 μl around blue pellet. Add 1ml of ice cold 80% ethanol (do not re-suspend) and spin samples for additional 10 mins at 4°C and >18,000x g.
3. Carefully remove all supernatant. Use a P10 pipette tip for last few μl of supernatant around pellet.
4. Air dry pellet at room temperature for 5 mins with lid opened.
5. Re-suspend in 5 μl nuclease free water and transfer to PCR tube (see **Note 19**, Figure 2B).
6. Add 1 μl of 10 mM dNTPs and 0.5 μl of 0.5 pmol/μl RT# primer to each each sample (see **Note 20**).
7. Denature RNA using the following thermocycler program:

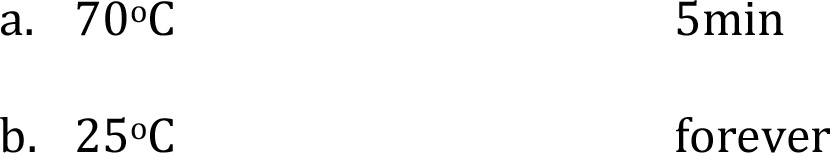
8. Add 13 μl of freshly prepared reverse transcription mix to each sample and mix by pipetting. Carry out reverse transcription using the following settings:

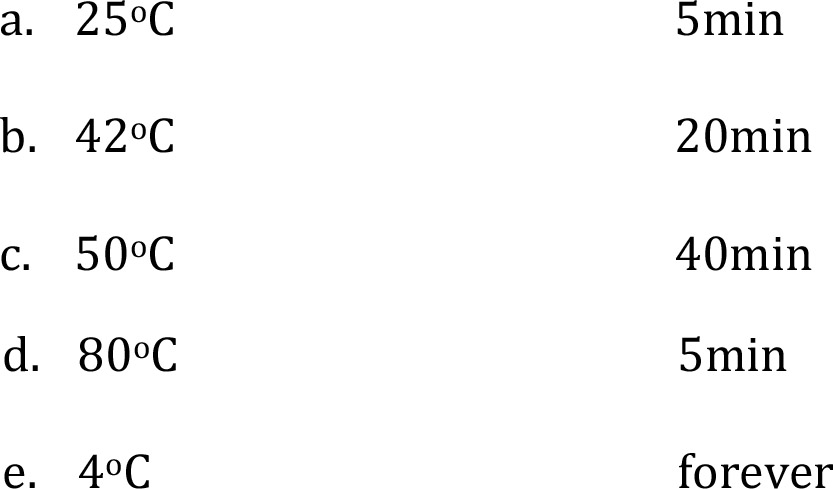
9. Add 1.65 μl of 1M NaOH to each sample and incubate at 98°C for 20min in order to cause alkaline hydrolysis of RNA that can interfere with the circligase reaction.
10. Neutralise samples by adding 20 μl of HEPES pH 7.3.
11. **Optional: At this stage samples to be multiplexed can be combined to reduce the number of samples to work with. It is recommended that negative controls are kept separate from working samples.**
12. Add 350 μl TE buffer, 0.75 μl Glycoblue, 40 μl 3 sodium acetate pH 5.5 and mix. Add 1ml of ice-cold 100% ethanol, mix again, then precipitate overnight at −20 °C.

### 3.9 cDNA purification and size selection

1. Spin samples for 20 mins at 4°C and >18,000x g.
2. Remove supernatant leaving ~50 μl around blue pellet. Add 1ml of ice cold 80% ethanol (do not re-suspend) and spin samples for additional 10 min at 4°C and >18,000x g.
3. Carefully remove all supernatant. Use a P10 pipette tip for last few μl of supernatant around pellet.
4. Air dry pellet at room temperature for 5 mins with lid opened.
5. Re-suspend pellet in 12 μl of 1x TBE-UREA loading buffer prepared using nuclease free water. Also add 6 μl of 2x TBE-UREA to 6 μl of a 1:30 dilution of low molecular weight marker.
6. Heat samples for 80 °C for 5 mins directly before gel loading.
7. Assemble the XCell III gel system according to manufacturers instructions using a 6% TBE-UREA gel (see **Note 21**) and fill chambers with TBE buffer.
8. Whilst samples are heating flush UREA out of gel wells
9. Load 12 μl of samples per well, allowing 1 well gaps between samples to facilitate gel cutting. Load ladder into a well at one end of the gel.
10. Run the gel for 40 min at 180V (see **Note 22**).
11. Whilst gel is running, prepare 0.5 ml microcentrifuge tubes for gel crushing. Carefully using a 16G syringe needle to pierce a clean hole in the bottom of 0.5 ml microcentrifuge tubes then inside new low-bind 1.5 ml microcentrifuge tubes.
12. Open the cassette and cut off the last lane containing the ladder. Stain the ladder using sybr safe and an appropriate imager, and use to construct a cutting guide that is printed out at 100% scale and wrapped in saran wrap (see **Note 23**). Alternatively, a pre-prepared cutting guide can be used (see **Note 22**, Figure 3).

**Figure 3:**
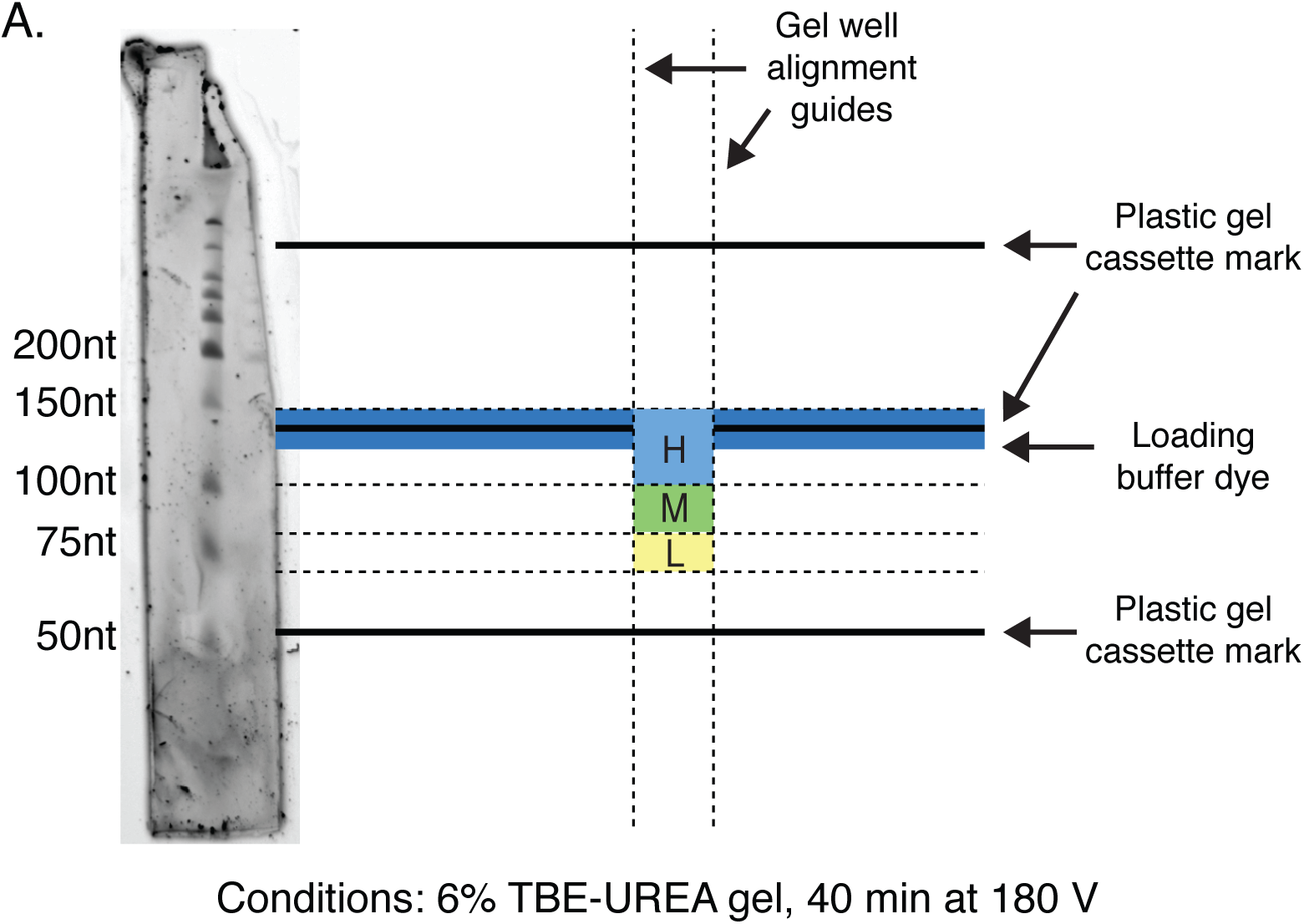
cDNA size selection. Example cutting mask determined from lowmolecular weight marker used for excising cDNA from denaturing TBE-UREA gel. Low, medium and high cut sites determined by denatured ladder are indicated, whilst gel cassette marks of the recommended TBE-UREA gel setup act as additional alignment guides.
13. Place the opened cassette upon the cutting guide and use markers to determine gel cut sites for each sample. The adapter and primer account for 52nt. Cut sizes and corresponding cDNA insert sizes are below (see **Note 24**):

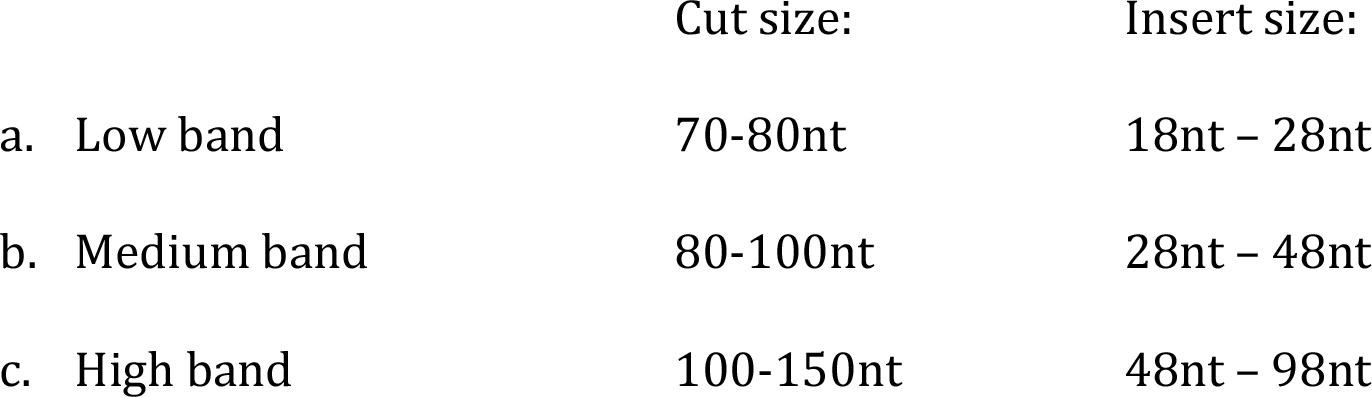
14. Transfer gel pieces to pierced 0.5 ml tubes sitting inside new low-binding 1.5 ml microcentrifuge tubes.
15. Spin samples at 10,000x g to crush gel pieces
16. Add 400 μl TE buffer and incubate for 1hr at 37°C shaking at 1100 rpm.
17. Place samples on dry ice to snap freeze samples ahead of rapid gel expansion.
18. Incubate samples for 1hr at 37°C shaking at 1100 rpm.
19. Whilst incubating, place 2 glass filters per sample into a costar spin-X column.
20. Cut end of P1000 tip to allow suction of gel pieces. Transfer buffer and gel pieces to prepared coster spin-X column placed in low-bind 1.5 ml microcentrifuge tube and spin at >18,000x g for 1min.
21. Transfer flow-through to Phase Lock Heavy gel columns together with 400 μl of neutral phenol-chloroform.
22. Incubate for 5 mins at 30 °C whilst shaking at 1100 rpm.
23. Separate phases by centrifuging at >18,000x g at room temperature for 5 mins.
24. Taking care not to touch the gel matrix, transfer the aqueous upper phase to a new low-binding 1.5 ml microcentrifuge tube.
25. Spin at >18,000x g for 1 min then transfer to a new low-binding 1.5 ml microcentrifuge tube.
26. Purify cDNA by adding 0.75 μl of Glycoblue and 40 μl of 3M sodium acetate pH 5.5 then mixing. Add 1ml of ice-cold 100% ethanol, mix again, then precipitate overnight at −20°C.

### 3.10 cDNA re-structuring

1. Spin samples for 20 mins at 4°C and >18,000x g.
2. Remove supernatant leaving ~50 μl around blue pellet. Add 1ml of ice cold 80% ethanol (do not re-suspend) and spin samples for additional 10 mins at 4°C and >18,000x g.
3. Carefully remove all supernatant. Use a P10 pipette tip for last few μl of supernatant around pellet.
4. Air dry pellet at room temperature for 5 mins with lid opened.
5. Resuspend cDNA pellet in 8 μl of freshly prepared circularisation mix and transfer to 0.2 ml PCR tubes.
6. Incubate for 1hr at 60 °C.
7. Add 30 μl of freshly prepared cut oligo mix to each PCR tube
8. Anneal the cut oligo with the following program:

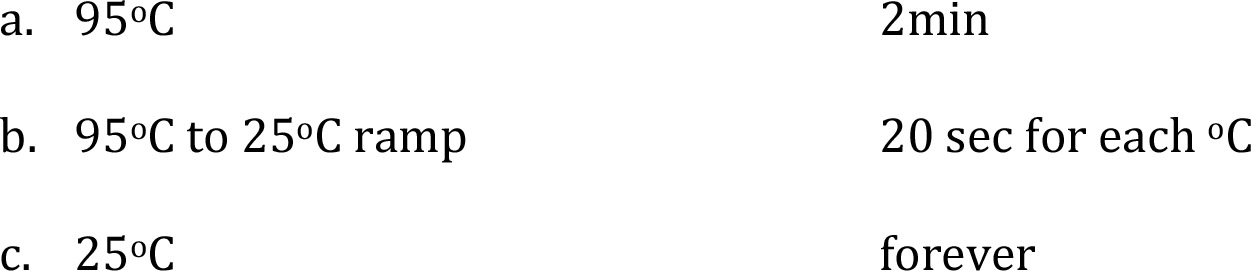
9. Add 2 μl of BamH1 to each PCR tube and incubate with the following program:

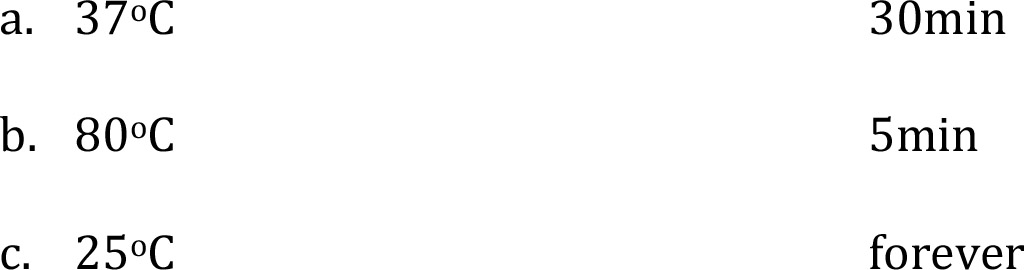
10. Transfer samples to new low-bind 1.5 ml microcentrifuge tubes.
11. Add 350 μl TE buffer, 0.75 μl Glycoblue, 40 μl 3 sodium acetate pH 5.5 and mix. Add 1ml of ice-cold 100% ethanol, mix again then precipitate overnight at −20°C.

### 3.11 cDNA library PCR

1. Spin samples for 20 mins at 4°C and >18,000x g.
2. Remove supernatant leaving ~50 μl around blue pellet. Add 1ml of ice cold 80% Ethanol (do not re-suspend) and spin samples for additional 10 mins at 4°C and >18,000x g.
3. Carefully remove all supernatant. Use a P10 pipette tip for last few μl of supernatant around pellet.
4. Air dry pellet at room temperature for 5 mins with lid opened.
5. Re-suspend the cDNA pellet in 22 μl nuclease free water
6. Add 1 μl of resuspended cDNA to 9 μl of test-PCR mix.
7. Perform test PCR with the following program:

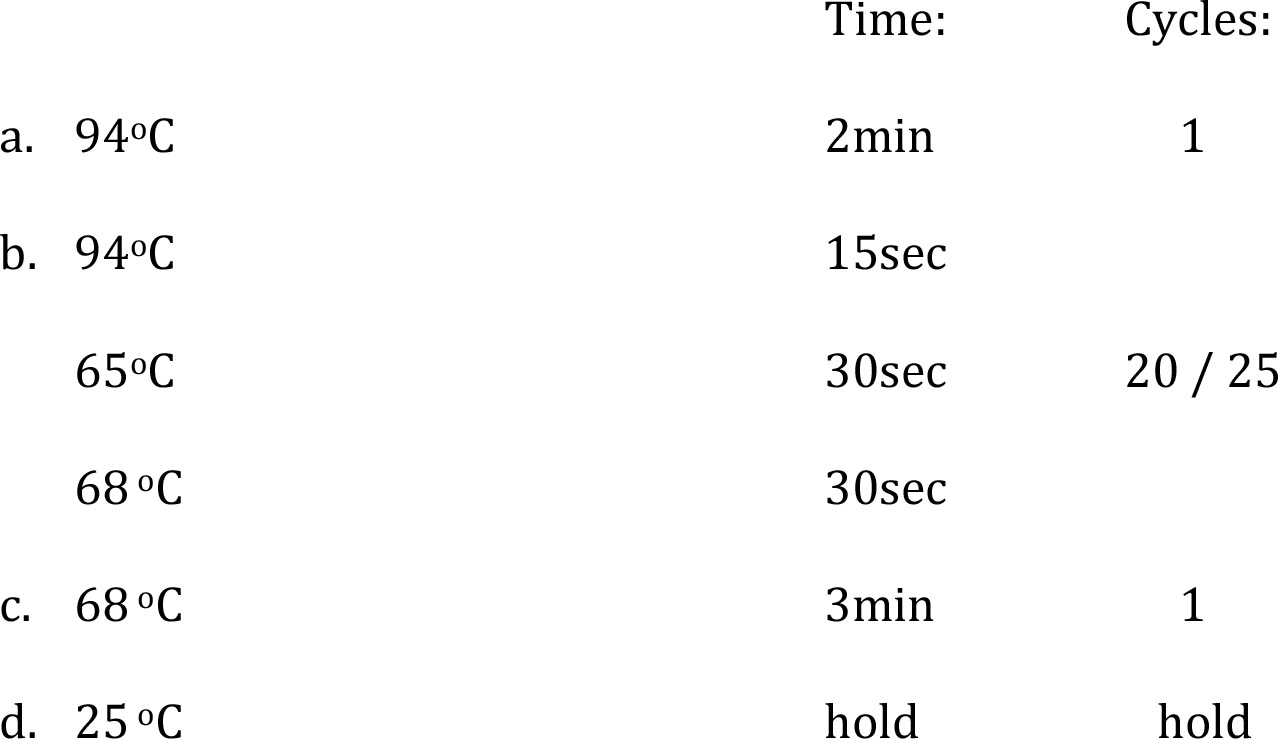 **Important: Following the PCR, do not open PCR tubes in same room where iCLIP library preparations (i.e. all steps before 3.11.8) were carried out. All post-PCR analysis and storage needs to be carried out in separate room to avoid amplicon contamination**.
8. Assemble the XCell III gel system according to manufacturers instructions using a 6% TBE gel and TBE running buffer.
9. Add 2 μl of 6x loading dye to each PCR sample and 10 μl of 1:30 low molecular weight marker. Load 12 μl of samples/ladder into gel wells.
10. Run gel for 30mins at 180V
11. Remove gel from cassette and stain for 5mins in SybrSafe in TBE buffer. Visualise using appropriate imager (Figure 4A). The P5/P3 primers account for 128nt of PCR product. Appropriate PCR band sizes are therefore:

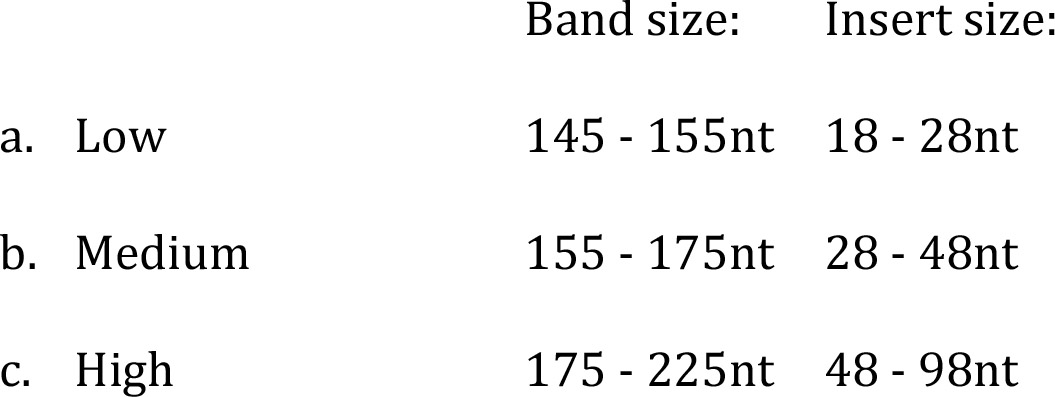

**Figure 4:**
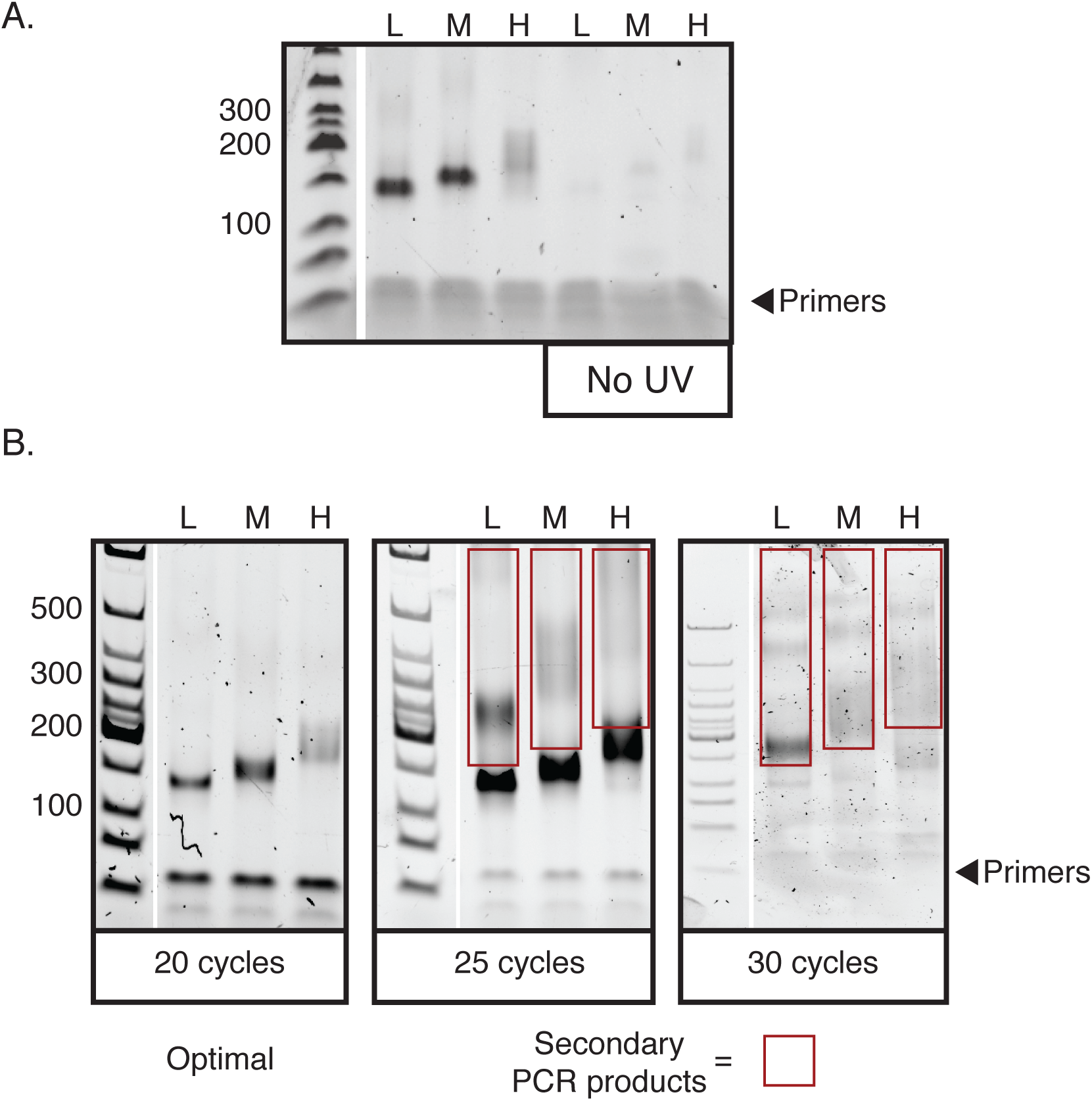
cDNA library PCRs. **A)** Example of final library PCR (3.11.13 – 3.11.17) showing low, medium and high bands. Both a low RNase sample and a no UV control sample are shown. **B)** Example of a cDNA library amplified with different PCR cycle numbers. Red boxes indicate secondary spurious products that migrate at a higher molecular weight to the expected products. **Optional: Steps 3.11.6-3.11.11 can be repeated with adjusted cycling Conditions to optimise cDNA library amplification (see Notes 25-26)**
12. Once optimal PCR conditions are determined, add 10 μl of resuspended cDNA to 30 μl of final-PCR mix:
13. Perform final PCR with the following program using one cycle less than optimal test-PCR (see **Note 27**):

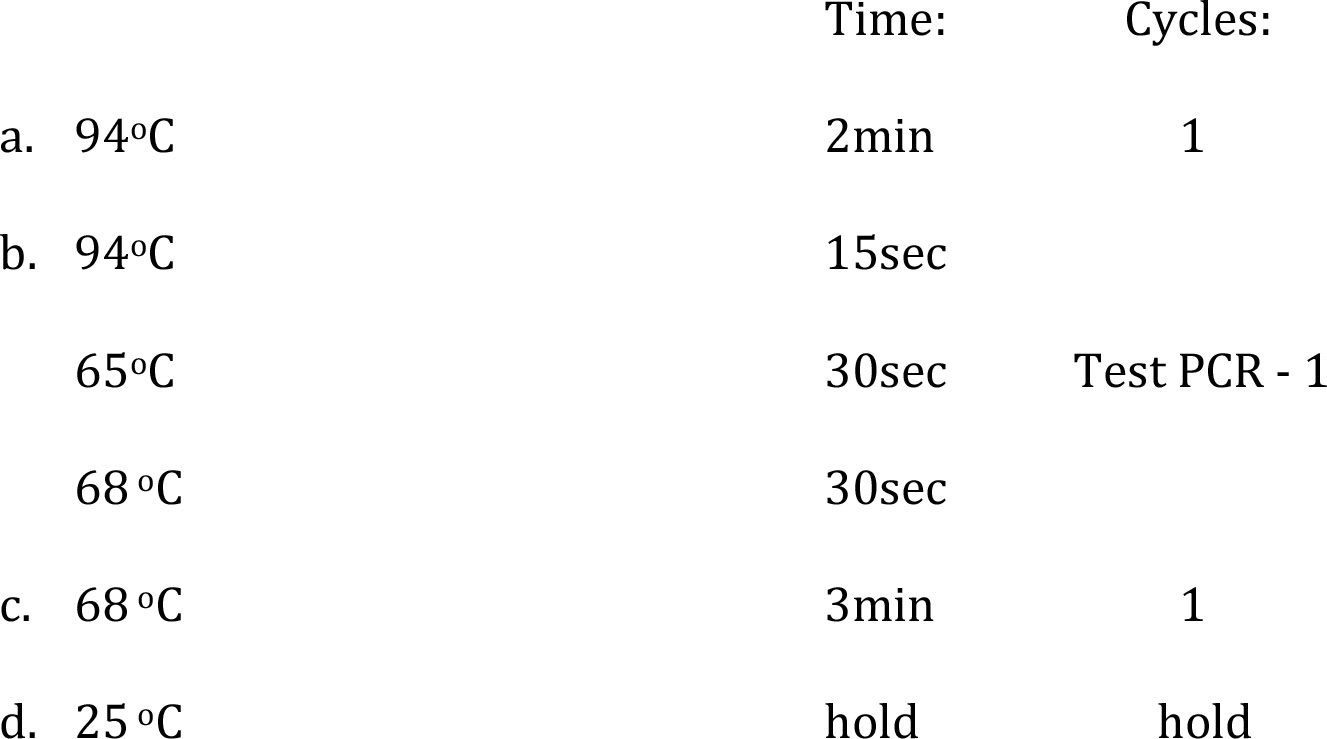
14. Assemble the XCell III gel system according to manufacturers instructions using a 6% TBE gel and TBE running buffer.
15. Add 2 μl of 6x loading dye to 10 μl aliquots of each final PCR sample and 10 μl of 1:30 low molecular weight marker. Load 12 μl of samples/ladder into gel wells.
16. Run gel for 30mins at 180V
17. Remove gel from cassette and stain for 5mins in SybrSafe in TBE buffer. Visualise using appropriate imager. **Optional: If sample is under-amplified then remaining PCR samples can be returned to thermocycler for 1-2 additional cycles before proceeding**.
18. Once optimal final PCR cycling number is confirmed then repeat final PCR (i.e. steps 3.11.13 – 3.11.14) using appropriate cycle number. Pool final PCRs together.
19. If PCR products are of correct size then aliquots of different sized PCR products can be combined in ratio of 1:5:5 (Low:Medium:High), or 1:1 (Medium:High) if no low band is included.

### 3.12 Library quantification

1. Prepare a 1:10 dilution of the final library, and then make serial dilutions to obtain 1:100, 1:1000 and 1:10,000 samples.
2. Prepare qPCR library quantification standards and non-template controls as per manufacturer guidelines.
3. In technical triplicate, add 16 μl of universal qPCR Master mix per qPCR plate/strip well.
4. Dispense 4 μl of non-template control, standards and serial library dilutions into reaction wells accordingly.
5. Seal plate, spin down and transfer to qPCR instrument. Run qPCR with the following cycling protocol:

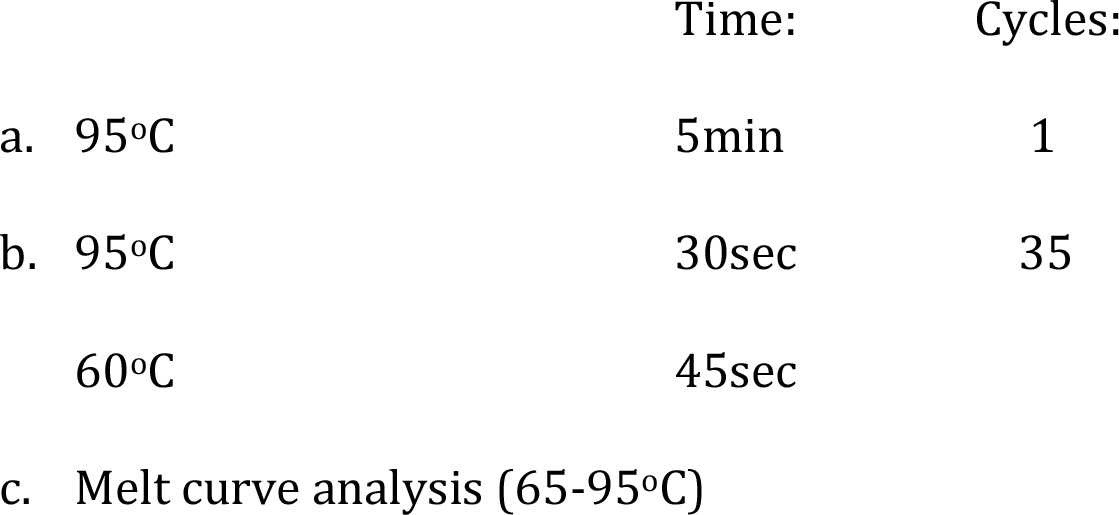
6. Determine concentration of library from qPCR standards and then correct for library type and insert size using pre-determined factor (see **Notes 28 and 29**).
7. Dilute library to 10nM and submit >10 μl to sequencing facility for sequencing. Provide facility with details about correction factor from step 3.12.6 so that it can be factored into any repeat quantification. Sequencing can be carried out in single-end 50-nucleotide runs (see **Notes 30 and 31**).

## 4. Notes

1. The adapter described in this protocol has the Li-Cor IRDye 800CW dye conjugated. An imager capable of detecting the 794nm wavelength is required. The high wavelength reduces background and increases sensitivity to levels that are required in iCLIP. It is expected that additional fluorophores with high wavelength characteristics may be viable alternatives with appropriate imagers e.g. > 600 nm.
2. It is critical that the pH is stably maintained during gel electrophoresis in order to prevent alkaline hydrolysis of the RNA. A pour-your-own gel is not suitable, whilst the Novex NuPage gels and system are strongly recommended when using the MOPS-SDS running buffer.
3. The use of 1/3^rd^ of a 10cm dish for each sample is a guideline suitable for standard cell lines. Combining pellets, or using smaller/larger vessels can increase sample size for hard to iCLIP proteins. It is also recommended that inputs are normalised by carrying out protein quantification following lysis and diluting to standard amounts.
4. Silencing of the RBP-of-interest should be confirmed through both qRT-PCR and western blotting to ensure specificity of knockdown. Appropriate silencing will reduce intensity of protein-RNA smear in step 3.6.13 rather than reducing size of smear.
5. It is important to keep the crosslinking conditions identical between experiments. This includes the height between UV bulbs and exposed samples. Finding a suitable tray that fits inside the crosslinker without interfering with the automatic cut-off switch will facilitate this.
6. The no UV control is critical to confirm that the signal is UV dependent and derived from cross-linked complexes. This control requires samples to be prepared in identical manner to experimental samples with the exception of UV cross-linking. Samples can be carried through to final PCR stage to assess library preparation efficiency and assess background levels of contamination contributing to library signal. In some cases where the RBP is both abundant and a particularly strong binder of RNA the no UV control may reveal some signal at the expected molecular weight. However, this should appear much weaker relative to the low RNase conditions (Figure 2A).
7. Less variable pellet sizes can be achieved by transferring 6 ml cell suspensions to a 15ml falcon tube, mixing by pipetting, then separating cells into 2 ml aliquots.
8. Protein G beads work well with most antibodies except sometimes rabbit. Protein A may work better with rabbit antibodies but protein A beads tend to stick to microcentrifuge tubes.
9. Wash steps are in 900 μl of indicated buffer unless specified. Washing steps involved magnetic separation of beads, supernatant removal then resuspension of beads in new buffer.
10. The no antibody negative control can use half the input to a normal sample and be carried through to final PCR stage as in case of the no UV control (see **Note 6**)
11. A positive control, such as anti-hnRNP C antibody [13,19,28], may be used to confirm that iCLIP is working in the hands of the experimenter, and provide a comparator of antigen size against RBP-of-interest. It is strongly recommended that a new iCLIP setup is first tested with a positive control antibody before moving on to a new RBP.
12. The integrity of the iCLIP protocol and the subsequent bioinformatics analysis strongly depends on optimal RNase digestion conditions [20]. In order to ensure samples aren’t over or under digested, it is critical that each new batch of RNase I is tested for activity on each batch of sample preparations used. In test experiments, dilutions of RNase I should span from 1:10 – 1:2000 (Figure 2A). In addition to fluorescent probe analysis following SDS-PAGE and nitrocellulose transfer, small aliquots of RNA extracted following proteinase-K digestions should be run on denaturing UREA gels in order to accurately assess RNA sizes corresponding to each digestion. Suitable digestions will produce an enrichment of RNA fragments in the size range of 50-300nt (Figure 2B).
13. A high RNase is used to confirm that the signal is sensitive to availability of RNase. It may also identify contaminating sources of signal or dimer/trimer complexes. Half the input to a normal sample should be used for the high RNase sample. High RNase will lead to a less heterogeneous pool of RNA lengths that will cause an intense signal above Mw of crosslinked antigen.
14. Enzymatic steps in section 3.5 and gel loading step in section 3.6 involve small volumes of 20 μl. Care should be taken to remove all carry over of wash buffers that can increase this volume. This can be achieved by removing the microcentrifuge tubes from the magnet for 15 seconds to allow beads to settle, then returning microcentrifuge tubes to the magnet and removing excess wash buffer.
15. Un-ligated adapter carried through into library preparation steps can be processed and lead to sample contamination. Various steps are carried out to ensure un-ligated adapter is removed. This includes stringent washing, transferring of washes to new tubes, nitrocellulose transfer following SDS-PAGE, and size-selection following cDNA synthesis.
16. The use of the antioxidant and reducing reagent maintains proteins in a reduced state during gel electrophoresis. Although use is optional, it is strongly recommended if the antibody bands run at a similar Mw to the RBP-of-interest. Without antioxidant and reducing reagent this will cause a band of reduced signal intensity across the protein-RNA complex smear in step 3.6.13
17. Cross-linking efficiency is ~1%. The Mw of the RBP-of-interest is therefore excluded since this position will include the majority of immunoprecipitated protein with no cross-linked and labelled RNA. Exclusion avoids potential saturation of proteinase K. Note that each extra 20nt of RNA will add ~7kDa to the molecular weight of the protein-RNA complex. Accordingly, high molecular weight RBPs or higher order complexes (e.g. dimers in Figure 2A) may benefit from being run for extended periods on higher percentage pre-cast gels in order to provide better resolution at these molecular weights.
18. The high RNase sample does not need to be excised. However, it is recommended that the no antibody and no UV negative controls are excised in order to assess library preparation integrity.
19. At this stage an aliquot of RNA can be run on a TBE-UREA denaturing gel to assess the size of the RNA fragments that have been extracted. The optimal size of RNA is between 50-300nt (Figure 2B). Since the cross-link site will be located at variable positions within RNA fragments, this size range leads to optimal cDNA insert sizes between 25-150nt.
20. Variation will exist between different batches of reverse transcription primer [13]. It is therefore recommended to test each new batch of reverse transcription primers to identify those that are producing optimal libraries. This can be achieved by taking a single low RNase sample up to step 3.8.5, then splitting the sample into equal aliquots for each reverse transcription primer. The remainder of the iCLIP protocol should be completed up to the test PCR stage (step 3.11.11) for each sample with no multiplexing. This will allow evaluation of the performance of each RT primer against one another under identical conditions.
21. TBE-UREA gels are required to fully denature cDNA for accurate size determination. Pre-cast TBE-UREA gels have short shelf-lives and should be not used outside of indicated dates.
22. Consistent conditions for cDNA gels allow the user to decide whether to use a previously made cutting guide or generate a new guide each time. Once it has been established that the denaturing conditions used can fully separate the low molecular weight marker into single stranded nucleotides (i.e. no ladder double bands due to partial denaturing), then it is possible to use a previous cutting guide if conditions have been kept identical (Figure 3).
23. Ladder staining should be carried out as quickly as possible as the TBE-UREA gels will start to swell once the cassette is open. A practice run on a ladder is recommended before the first iCLIP experiment, whilst a pre- 27 made cutting guide can also be used as indicated in **Note 22**.
24. Three bands are cut from each sample. This is to avoid PCR bias towards shorter fragments at later steps. The lowest band contains short insert sizes that are less mapable than the medium and high bands. However, they may contain short RNAs including miRNAs. Unless short RNAs are desirable, the crushed low band can be stored at −20°C for processing at a later date if required. Should PCR analysis reveal inaccurate and high cutting, then the stored low band may also be processed to produce optimal cDNA libraries.
25. Over amplification of cDNA can lead to secondary products appearing above the expected size (Figure 4B) [13]. At higher cycle numbers these products are too large to migrate on the 6% TBE gels. These secondary products carry the necessary sequences required for flow-cell hybridisation and so should be removed through PCR optimisation.
26. A low cycle final PCR (~15-21 cycles) will produce well-diversified libraries, whilst high PCR cycles (> 25 cycles) will be produce libraries where limited products are sequenced many times over. It is suggested that an initial test PCR of 25 cycles is carried out to ascertain where the amplification stands relative to Figure 4B libraries. Appropriate cycle number changes can then be made for a second test PCR to finalise conditions. For example, if the test PCR has a similar appearance to the 25 PCR cycles in Figure 4B then the next test PCR should be reduced by 4-5 cycles in order to get optimal amplification at 20-21 cycles.
27. The final PCR uses 2.5 fold more concentrated input cDNA than the test PCR. Accordingly, an adjustment of 1 reduced cycle results in optimal final PCR cycling number.
28. In our hands we find that the calculated concentration divided by 2 results in accurate values for illumina sequencing. However, it is recommended that relationships between qPCR and sequencing cluster density are closely monitored over initial sequencing runs in order to optimise in a new iCLIP setup.
29. Alternatives to qPCR quantification can include Agilent Bioanylser/ TapeStation analysis together with Qubit quantification e.g. [29].
30. Following high throughput sequencing the bioinformatics analysis of iCLIP libraries requires de-multiplexing, mapping, collapsing PCR duplicates using random barcodes and determination of the crosslink site. Note that the nucleotide immediately preceding the first nucleotide of a mapped read is considered the site of crosslinking. After replicate comparisons, biological and technical replicates may be merged before subsequent analysis of the RBPs binding landscape.
31. Following bioinformatics removal of PCR duplicates based on random barcode evaluation and the merging of technical/biological replicates, an iCLIP library will ideally have ~1 × 10^6^ unique reads for analysis. However, libraries with >1 × 10^5^ can allow sufficient information to perform a basic analysis.

## 5. Acknowledgements

The iCLIP protocol described is based on previous versions developed in the Ule and Konig labs by many individuals. I would like to extend thanks to all those who have contributed to the methods development over the years. I would also like to thank Prof. Jernej Ule for critical reading of the chapter, and Prof. Jernej and Flora Lee for providing the adapter oligo used in this protocol. This work is supported by an Edmond Lily Safra fellowship to C.R.S.

